# A deep learning framework for imputing missing values in genomic data

**DOI:** 10.1101/406066

**Authors:** Yeping Lina Qiu, Hong Zheng, Olivier Gevaert

## Abstract

**Motivation:** The presence of missing values is a frequent problem encountered in genomic data analysis. Lost data can be an obstacle to downstream analyses that require complete data matrices. State-of-the-art imputation techniques including Singular Value Decomposition (SVD) and K-Nearest Neighbors (KNN) based methods usually achieve good performances, but are computationally expensive especially for large datasets such as those involved in pan-cancer analysis.

**Results:** This study describes a new method: a denoising autoencoder with partial loss (DAPL) as a deep learning based alternative for data imputation. Results on pan-cancer gene expression data and DNA methylation data from over 11,000 samples demonstrate significant improvement over standard denoising autoencoder for both data missing-at-random cases with a range of missing percentages, and missing-not-at-random cases based on expression level and GC-content. We discuss the advantages of DAPL over traditional imputation methods and show that it achieves comparable or better performance with less computational burden.

**Availability:** https://github.com/gevaertlab/DAPL

## Introduction

The Cancer Genome Atlas (TCGA) project has provided researchers a rich database to study the molecular basis of cancer (Cancer Genome Atlas Research, et al., 2013). Pan-cancer analyses enable a better understanding of the genomic commonalities and differences across tumor types and subtypes, which makes it possible to extend diagnosis and treatment from one cancer type to another with a similar genomic profile (Cline, et al., 2013). RNA sequencing data and DNA methylation data have been widely investigated in pan-cancer analyses to identify cancer driver genes or biomarkers (Byron, et al., 2016; Gevaert, et al., 2015; Kulis and Esteller, 2010; Litovkin, et al., 2015; Tomczak, et al., 2015). Many such studies on pan-cancer genomics require complete datasets (Champion, et al., 2018). However, missing values are frequently present in these data due to various reasons including low resolution, missing probes, and artifacts (Baghfalaki, et al., 2016; Libbrecht and Noble, 2015). Therefore, practical methods to handle missing data in pan-cancer genomic datasets are needed for effective downstream analyses.

One way to complete the data matrices is to ignore missing values by removing the entire feature if any of the samples has a missing value in that feature, but this is usually not a good strategy as the feature may contain useful information for other samples. The most preferable way to handle missing data is to impute their values in the pre-processing step. Many approaches have been proposed for this purpose, including replacement using average values, estimation using weighted K-nearest neighbor (KNN) method (Faisal and Tutz, 2017; Troyanskaya, et al., 2001), and estimation using singular value decomposition (SVD) based methods (Troyanskaya, et al., 2001). In recent years, a branch of machine learning which emerged based on big data and deep artificial neural network architectures, usually referred to as deep learning, has advanced rapidly and shown great potential for applications in bioinformatics (Min, et al., 2017). Deep learning has been applied in areas including genomics studies (Chen, et al., 2016; Leung, et al., 2014), biomedical imaging (Chen, et al., 2016), and biomedical signal processing (Wulsin, et al., 2011). Deep learning based methods have also been proposed to solve the missing data problems in various contexts and shown promising results (Beaulieu-Jones and Moore, 2017; Jaques, et al., 2018; Vincent, et al., 2008). Among the competitor algorithms for genomic data imputation, KNN and SVD are still considered the de facto standard because they can be easily implemented and have been reported to achieve best performances in missing value estimation problems (Aghdam, et al., 2017). However, when applied to very large datasets, KNN and SVD have severe drawbacks in computational overhead. Both methods are not model based and hence require significant computation at evaluation time, which makes them less practical in large-scale genomic data imputation.

In this study, we propose to leverage the denoising autoencoder (DAE) structure to predict missing values in pan-cancer genomic analysis. We demonstrate that by using a denoising autoencoder with partial loss (DAPL), it is possible to achieve significant improvement over a standard denoising autoencoder. We also show that the performance is on a par with the most state-of-the-art competitor algorithms while requiring less computational cost at evaluation time, for both data missing-at-random cases across a range of missing percentages, and missing-not-at-random cases based on expression level and GC-content.

## Materials and Methods

### Datasets

We used the pan-cancer RNA sequencing data and DNA methylation data from TCGA. We preprocessed the raw RNA sequencing data by removing the genes with NA values, and doing a log transformation followed by z-score transformation. The resulting matrix has 11069 samples and 17175 genes. We used the pan-cancer DNA methylation data to validate the algorithm on a larger and wider (feature dimension >> sample size) dataset. The validation data was processed in the same way as the RNA sequencing data, which resulted in a matrix with 9664 samples and 269273 features (CpG sites).

### Autoencoder

An autoencoder is an unsupervised deep neural network that is trained to reconstruct an input *X* cby learning a function *h_w,b_*(*X*) ≈ *X*. This is done by minimizing the loss function between the input *X* and the network’s output *X’: L* (*X, X*’). The most common loss function is the root mean squared error:

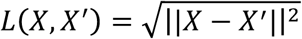

An autoencoder consists of an encoder and a decoder. The encoder transforms the input to a latent representation, often such that the latent representation is in a much smaller dimension than the input (Ballard, 1987). The decoder then maps the latent embedding to the reconstruction of *X*. An autoencoder is often used as a dimensional reduction technique to learn useful representation of data (Sakurada and Yairi, 2014).

A denoising autoencoder (DAE) aims to recover the noise-free, original input through deep networks given a noisy input (Vincent, et al., 2008). In each iteration of training, noise is added to the input *X* to obtain 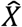. The loss is computed between the original *X* and the reconstructed 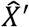.

### Denoising autoencoder with partial loss

When the added noise to DAE is in the form of masking noise, i.e., a random fraction of the input is set to 0, the DAE can be trained to estimate missing values (Vincent, et al., 2010). The 0s in the training data simulate the missing elements and the reconstructed noise-free data will fill those positions with estimated values. However, a standard DAE aims to reproduce the whole dataset without focusing on the missing elements, while some of the other state-of-the-art imputing methods, such as SVD, focus on learning the missing elements iteratively. As a result, the performance of a standard DAE in estimating missing values is not on par with SVD, or KNN based methods.

To ameliorate this problem, we implemented a denoising autoencoder with partial loss (DAPL) to focus the neural network’s learning on the missing positions. We minimized the root mean square error (RMSE) loss only on the missing values between the original input *X*, and the reconstructed 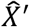:

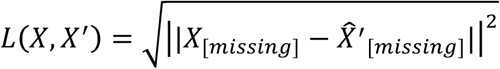

### Missing data simulations

With the TCGA RNA sequencing dataset, we only used the features that have values for every sample to construct a ground truth complete data. Then the complete data was split 60/15/25% for training, validation, and testing.

We simulated two missingness scenarios: at random and not at random. In the missing at random case, missing values in the testing data are in this form: a fixed fraction of elements at random positions were masked by setting their values to NA. We experimented with a range of missing fraction: 5%, 10%, 30%, 50% and 80%. We artificially introduced missingness to the complete training data by masking certain values. The autoencoder was initially trained such that the training data was masked to have the same fraction of random missingness as the testing data it was applied on. Then it was also trained on training data masked at different fraction of missingness than the testing data for comparisons. We then computed the RMSE error between the test ground truth and the imputed values. In the missing not at random case, we implemented two scenarios for missing. The first one was based on gene expression level (Conesa, et al., 2016). We tested on cases where a random half of the genes with expression level at the lowest 5% or 10% are missing. The autoencoder was trained on the same simulated scenario, where a random half of the genes with expression level at the lowest 5% or 10% were masked as missing during training. The second scenario was based on GC content, the percentage of nitrogenous bases on a RNA fragment that are either guanine(G) or cytosine(C). Too high or too low GC content influences RNA sequencing coverage, and potentially results in missing values (Chen, et al., 2013). We tested on cases where a random half of the genes with GC-content at the highest 5% or 10% are missing. Similarly, training was done on the same simulated scenarios.

### Evaluation methods

The model hyper parameters we tested include number of hidden layers, node size and dropout. We evaluated twelve autoencoder structures including three different number of layers (Figure 1, Table 1): 1) three layers with a layer-wise size reduction factor of 2, 4, 8 and 16-folds; 2) five layers with a layer-wise size reduction factor of 2, 3, 5, 8 and 10-folds; 3) seven layers with a layer-wise size reduction factor of 2, 3 and 4-folds. Layer-wise size reduction factor is the amount of reduction in the number of nodes in one layer of an autoencoder compared to its previous layer in the encoder portion. For example, a reduction factor of two means one layer has approximately half of the number of nodes than its previous layer. Due to the symmetric architecture of autoencoders, the reduction factor in the encoder is the same as the layer-wise increase factor in the decoder. We will refer to this number as the layer-wise reduction factor. Dropout is a regularization technique to reduce autoencoder overfitting (Srivastava, et al., 2014). At training stage, individual units were dropped out of the network with a probability called dropout rate.

**Table 1.** Autoencoder with Partial Loss (DAPL) structural variations

| Layer Number | Reduction Factor | Bottleneck Size |
| --- | --- | --- |
| 3 Layers | 2 | 8000 |
|  | 4 | 4000 |
|  | 8 | 2000 |
|  | 16 | 1000 |
| 5 Layers | 2 | 4500 |
|  | 3 | 2000 |
|  | 5 | 800 |
|  | 8 | 250 |
|  | 10 | 170 |
| 7 Layers | 2 | 800 |
|  | 3 | 400 |
|  | 4 | 200 |

**Figure 1.**
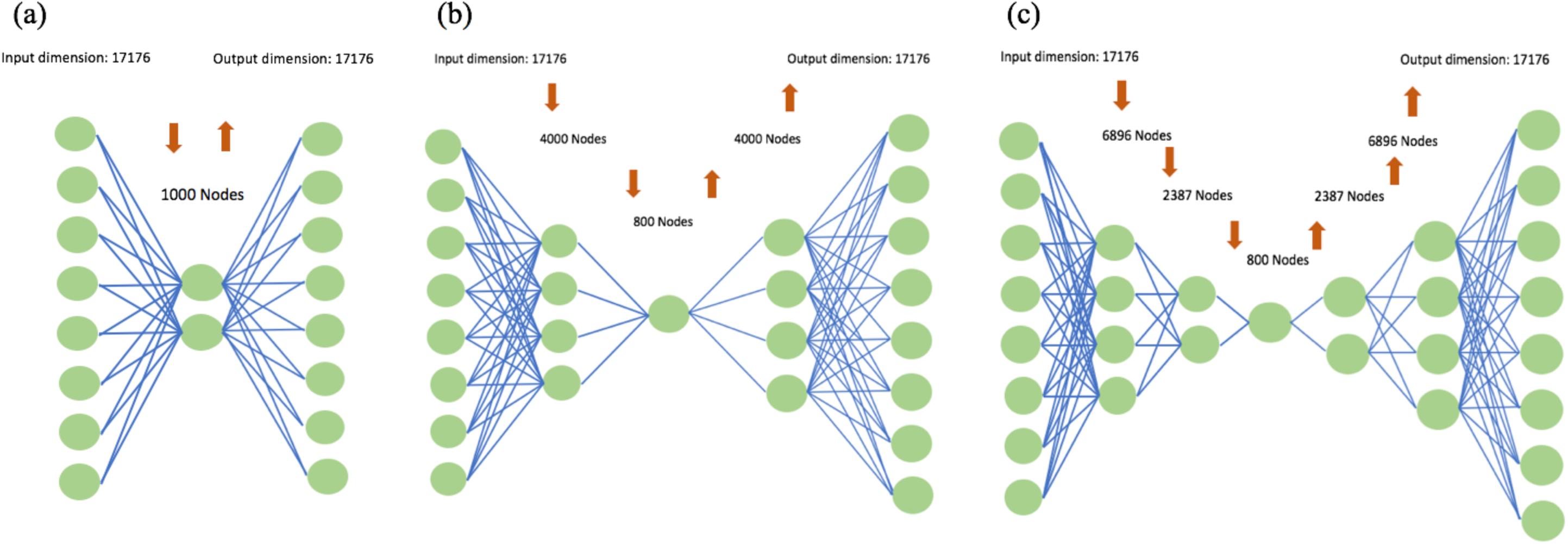
Schematics showing (a) a three layer autoencoder with bottleneck size of 1000. (b) a five layer autoencoders with bottleneck size of 800, and (c) a seven layer autoencoders autoencoder with bottleneck size of 800

To evaluate denoising autoencoder with partial loss, we compared it to the other most commonly used missing value estimation methods: KNN method, SVD based method, standard DAE without loss modification, and column average imputation. KNN selects K number of samples which are most similar to the target sample with a missing gene based on Euclidean distance, and which all have values present in that gene. Imputation is a weighted average of the values of that gene in those K samples. We chose K=10 in our evaluations based on a study which reported that K in the range of 10-25 gave the best imputation results (Troyanskaya, et al., 2001). SVD method decomposes the data matrix to the linear combination of eigengenes and corresponding coefficients. Genes are regressed against K most significant eigengenes, during which process the missing genes are not used (Hastie, et al., 1999). The obtained coefficients are linearly multiplied by eigengenes to get a reconstruction with missing genes filled. This process is repeated until the total change in the matrix reaches a certain threshold. The reconstruction performance of SVD depends on the number of eigengenes selected for regression, and we tested a range of values to determine the optimal K for SVD performance. Standard DAE was implemented as described previously, with loss defined over the entire data matrix. Column average imputation estimated the missing gene by taking the average of the gene in all samples where that gene was present.

We evaluated each test sample and computed RMSE error of the missing values in that sample

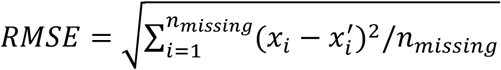

Where *x*_*i*_ is the element in the ground truth, and is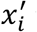is the element in the reconstructed data. Then we compared the RMSE distribution of the test samples for each of the method. Since SVD and KNN need to compute each time a new sample is tested, it is computationally expensive to evaluate all the test data. We randomly chose 100 samples from the test set for a fair comparison of all methods.

The method was validated on the TCGA pan-cancer DNA methylation data. Deep learning methods are usually faced with the challenges of severe overfitting and high-variance gradients when processing datasets that have a small sample size but high feature dimension (Liu, et al., 2017). We chose the methylation data where feature dimension is far greater than sample size to validate our algorithm’s robustness and applicability in real world problems. 5%, 30% and 80% random missing were tested on a five layers autoencoder which has a layer-wise size reduction of 100-fold with bottleneck size of 800. Since it requires large computation resource to perform competitor algorithms on this large dataset, we did not run analysis on other random missing percentages and missing not-at-random cases due to hardware limitations.

## Results

### DAPL outperforms DAE

We compared the proposed AE models for different missingness scenarios and also evaluated different autoencoder structures. First we tested missing at random cases at varying percentage (Figure 2). 5%, 10%, 30%, 50%, and 80% random elements in the data were masked respectively, and models were compared on the reconstruction root mean square errors (RMSE). Using a five hidden layers with 800 node bottleneck autoencoder structure, DAPL achieves significant improvement over the standard DAE for the comparisons between DAE and DAPL in all missing percentages (Wilcoxon rank sum test, P-value < 0.001).

**Figure 2.**
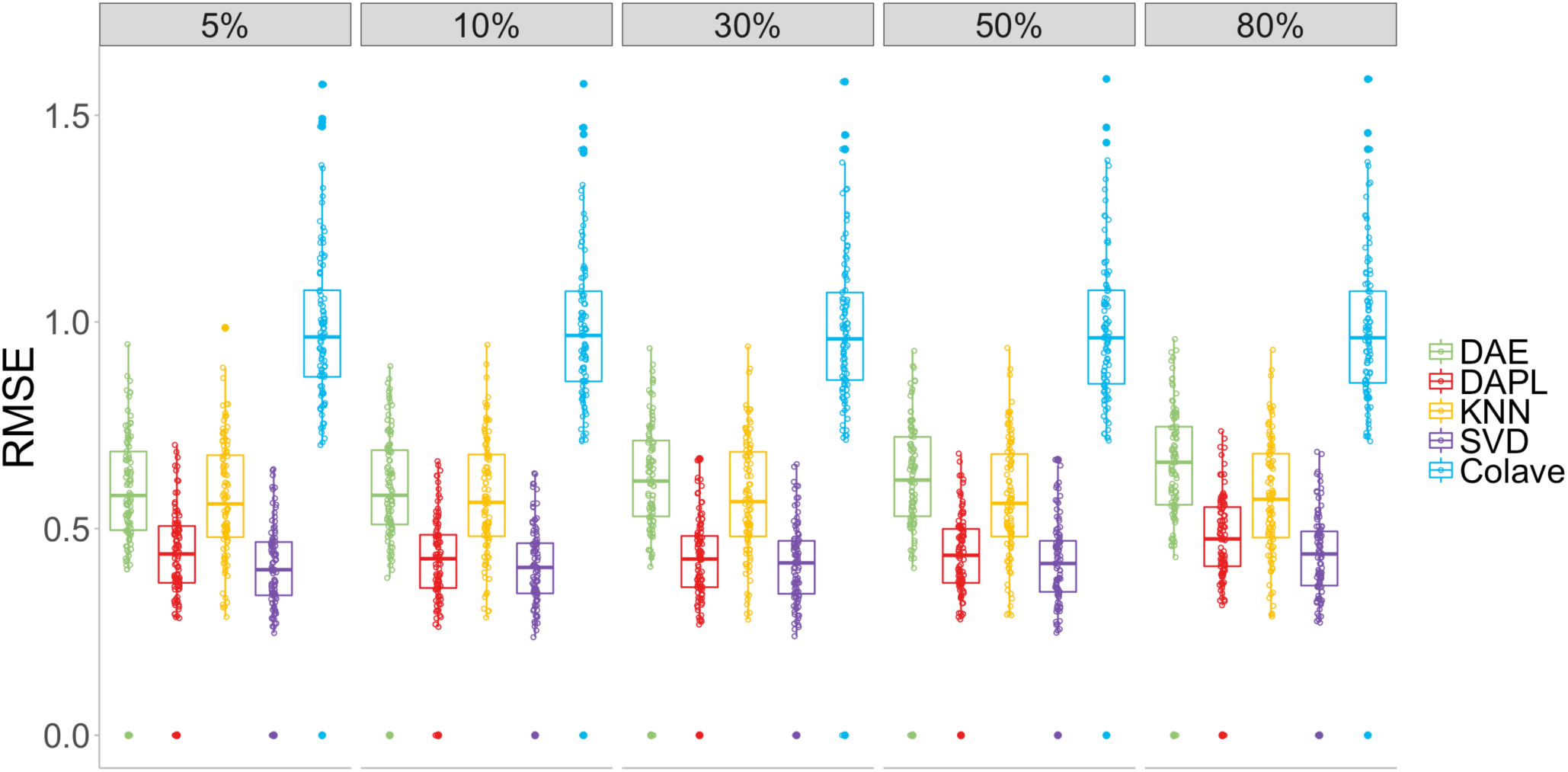
Comparisons of imputation methods measured in root mean square error (RMSE) for missing at random cases. From left to right: testing is done at 5%, 10%, 30%, 50%, and 80% random missingness. The method used from left to right: standard denoising autoencoder (DAE), denoising autoencoder with partial loss (DAPL), 10 nearest neighbor method (KNN), iterative singular value decomposition (SVD), and column average estimation (Colave).

To determine the optimal number of eigengenes K used for regression in the SVD method, we experimented with 1%, 2.5%, 5%, and 10% of the eigengenes and concluded that 2.5% - 5% gives the best imputation results. The comparisons were done with K=2.5% of eigengenes. DAPL reaches an average error very close to the best performing SVD. 10-KNN method gives worse average error than SVD and DAPL, but slightly better than a standard denoising autoencoder without loss modification. Column average results in worst imputation performance among all methods considered. The situation does not change with changing noise levels.

We tested several dropout rates and found that dropout applied to hidden layers did not improve imputation performance. We also tested a different way of masking at training stage: we randomly selected a fixed fraction of columns in the whole data set, and set the entire columns to zero. We found it did not improve the imputation performance of DAPL.

### Optimal training for missing at random

Next, we investigated the effect of different masking percentages during training of DAPL for predicting at specific missing levels. For each test case from the five missingness levels, DAPL was trained on all masking levels to find out which training scheme gives the best performance. Our results show that if training is done on the same simulated scenario as the testing case, i.e., the same level of missingness in testing is introduced during training, the performance is better than if training is done on a different scenario (Figure 3). However, it is also shown that when the testing missingness level is smaller than 50%, training at 30% masked achieved reasonable performance at all levels.

**Figure 3.**
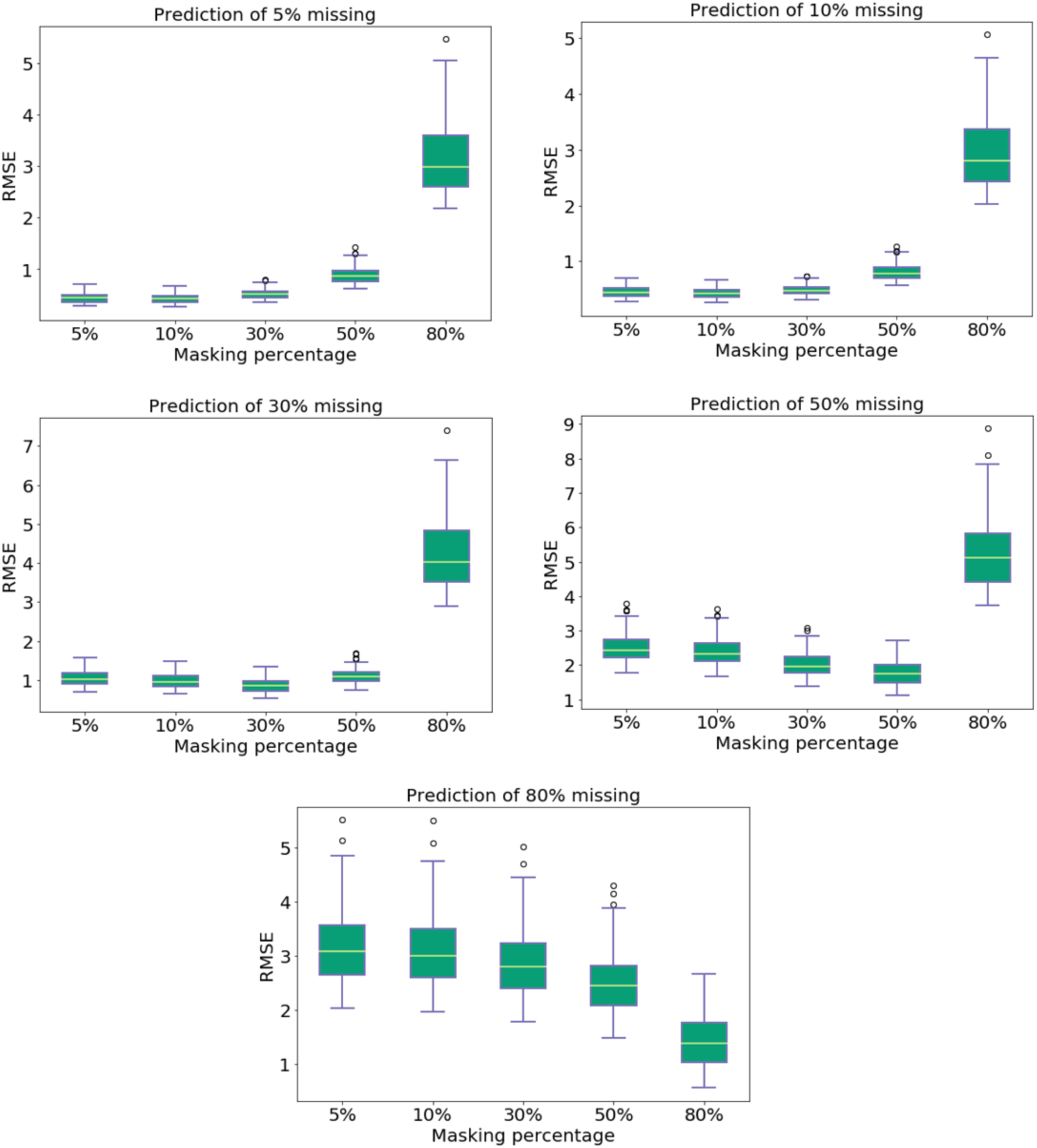
Effect of different masking percentages during training of denoising autoencoder with partial loss, measured in root mean square error (RMSE), for predicting at each of the 5 missing levels from top left to bottom right: 5%, 10%, 30%, 50%, 80% missing.

### Optimal architecture for DAPL

We evaluated twelve autoencoder architectures including three, five and seven layer networks with a range of layer-wise reduction factors. We found that reduction factor has a noticeable effect on imputation accuracies (Figure 4). The model complexity decreases with increasing layerwise reduction factor. In general, the trend is that the RMSE gets higher when the model becomes simpler. However, this is not strictly true. The RMSE trend drops slightly at the beginning for five layer autoencoders, and does not increase steeply within a certain range. This suggests that the optimal DAPL architecture is not the most complex one, and hence it is possible to reduce complexity without heavily degrading performance.

**Figure 4.**
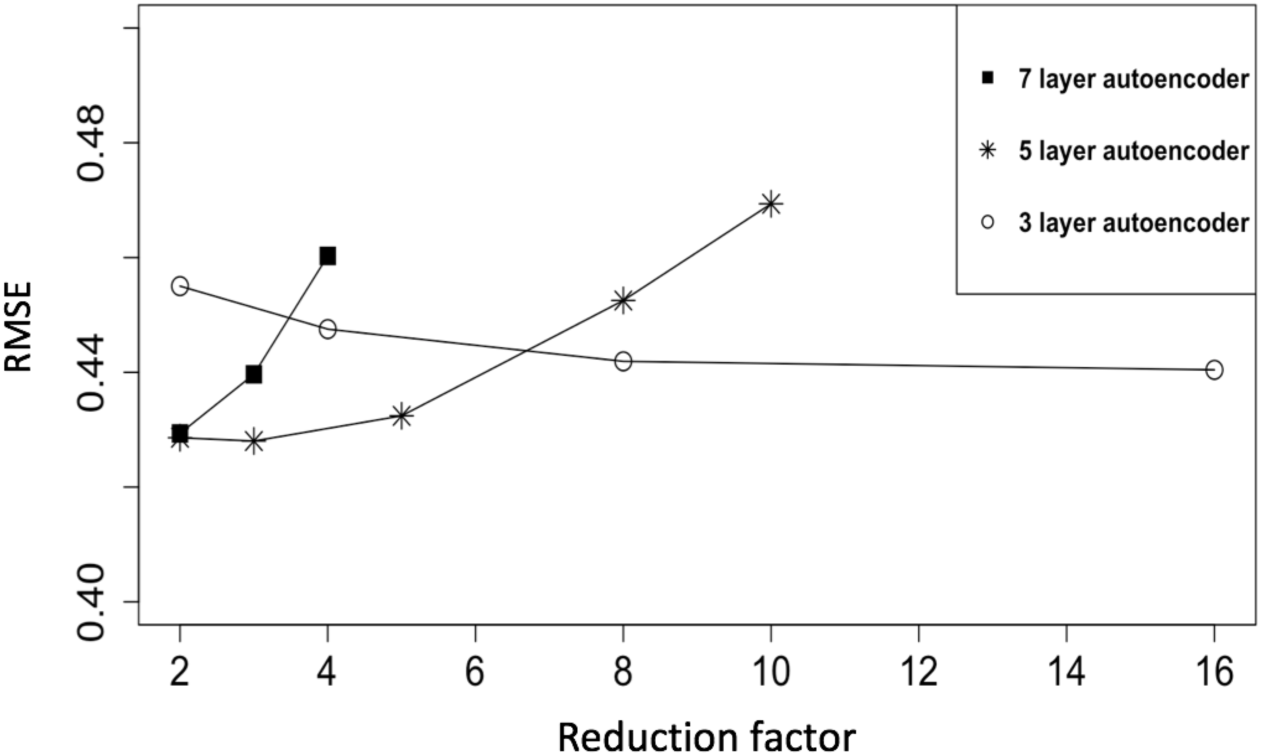
Results of autoencoder structural variations. Root mean square error (RMSE) is measured against layerwise size reduction factor for each of 3 layer, 5 layer, and 7 layer autoencoders.

### DAPL outperforms SVD for missing not at random

Missing not at random cases include missing based on expression level where a random half of the genes with expression level at the lowest 5% or 10% are missing, and missing based on GC content where a random half of the genes with GC-content at the highest 5% or 10% are missing. Both were performed with an autoencoder with five hidden layers and bottleneck size of 800. For missing based on expression level cases, DAPL outperforms SVD, achieving the lowest RMSE among all methods in both missing percentages (Figure 5). For missing based on GC-content cases, SVD has the lowest RMSE compared to other methods, followed by DAPL (Figure 5).

**Figure 5.**
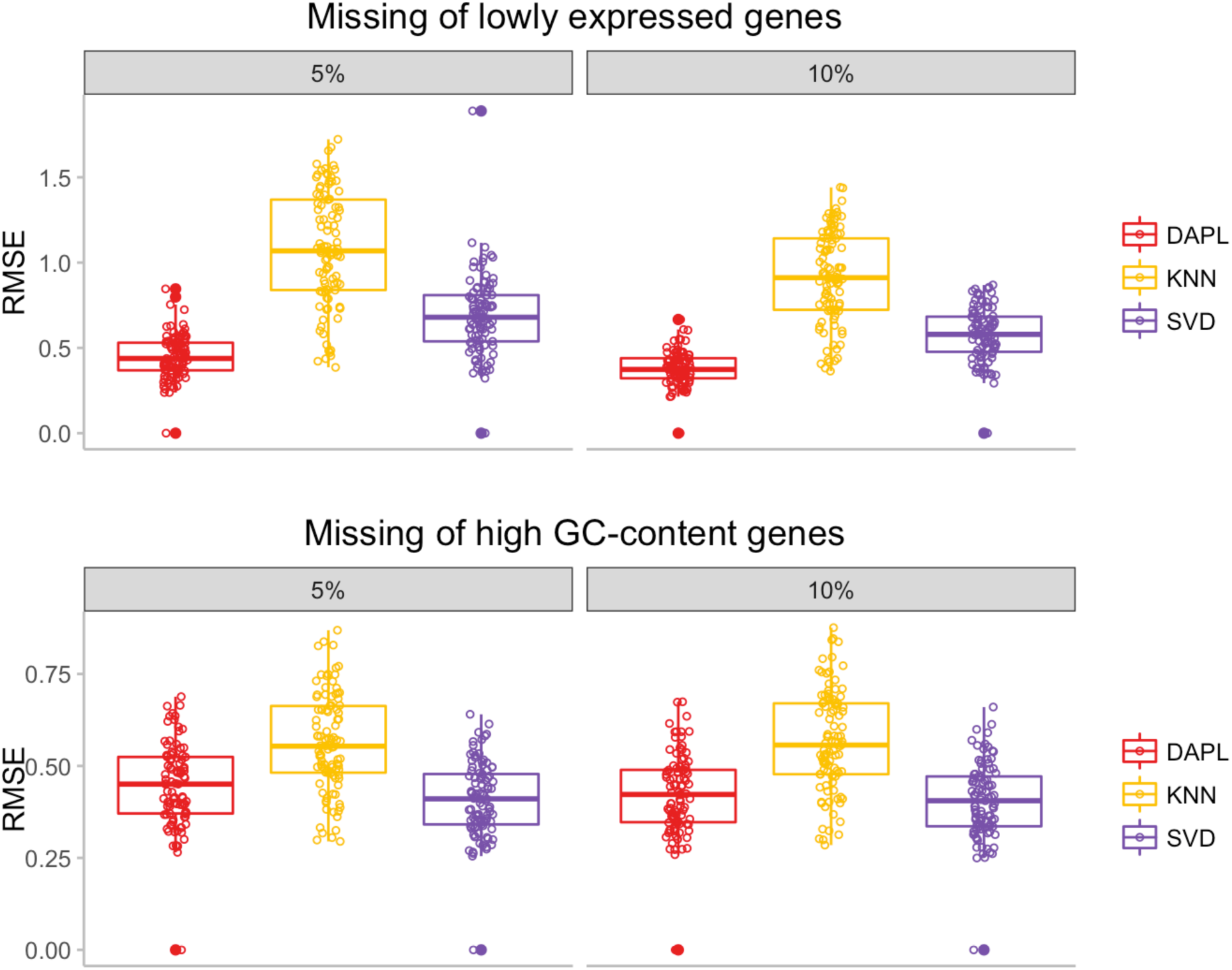
Comparisons of imputation methods measured in root mean square error (RMSE) for missing not at random cases. Top row from left to right: a random half of the genes with expression level at the lowest 5% and 10% are missing. Bottom row from left to right: a random half of the genes with GC-content at the highest 5% or 10% are missing. The method used from left to right: denoising autoencoder with partial loss (DAPL), 10 nearest neighbor method (KNN), and iterative singular value decomposition (SVD).

### DAPL method performs similarly on DNA methylation data

The method is validated on the TCGA pan-cancer DNA methylation data. This is to test whether DAPL is robust enough to work on data where the feature space is far greater than the sample size, which is a scenario in neural network-based methods that usually raises issues such as overfitting, and hence results in poor performances. On the methylation data, we see that due to increased data size, all of the imputation methods have higher RMSE than in the RNA sequencing data. On all simulation cases SVD still outperforms others, but DAPL shows similar or better result than KNN (Figure 6).

**Figure 6.**
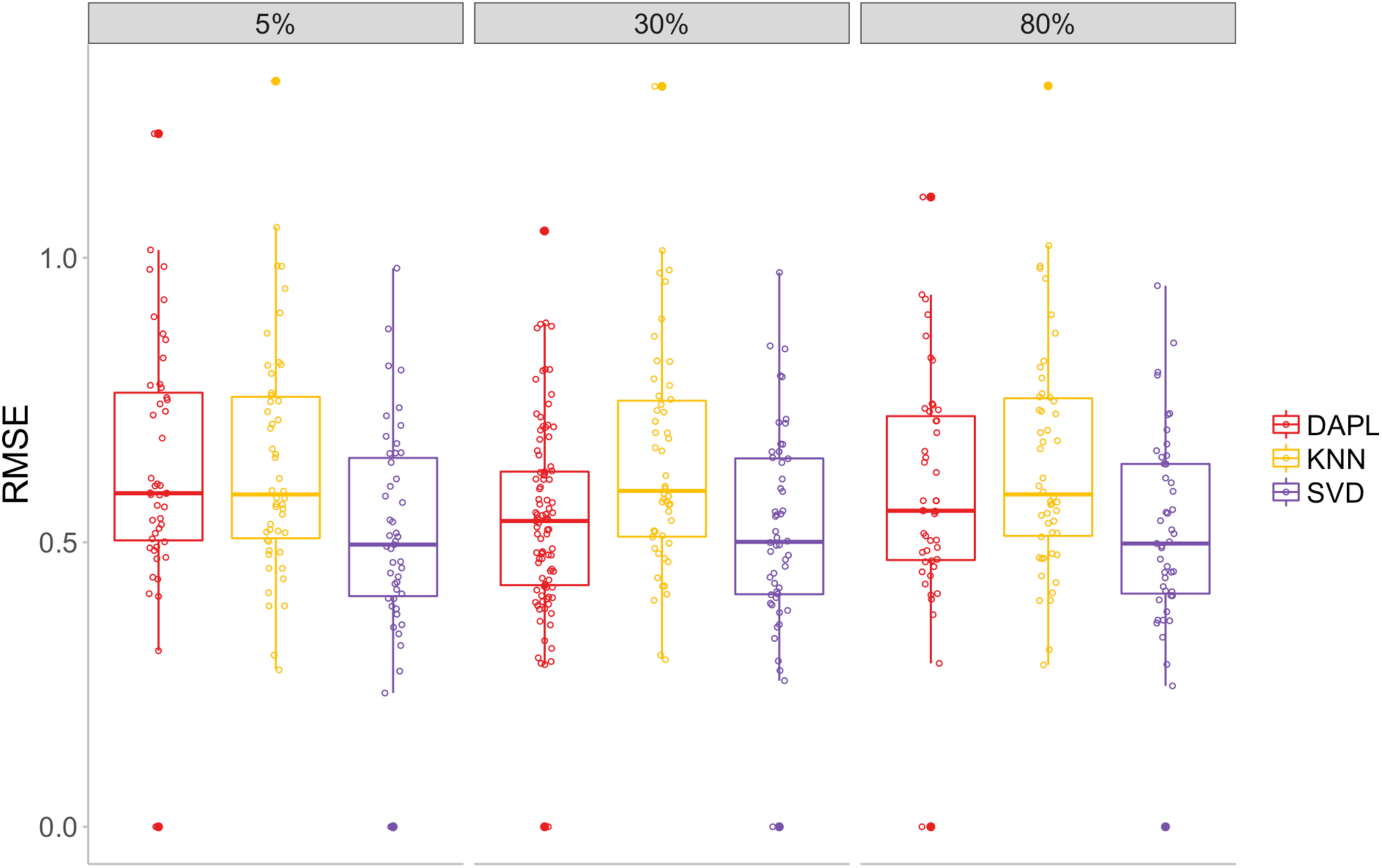
Comparisons of imputation methods measured in root mean square error (RMSE) for missing at random cases on DNA methylation data. From left to right: testing is done at 5%, 30%, and 80% random missing. The method used from left to right: denoising autoencoder with partial loss (DAPL), 10 nearest neighbor method (KNN), and iterative singular value decomposition (SVD).

### DAPL is the fastest method for evaluation

All computations were done on a 20 core cluster with Intel Xeon 2.40GHz CPUs. In terms of evaluation time, the KNN method took on average 8405 seconds to estimate missing values for a dataset with 100 samples and 16176 genes, with 5% of the entries missing, while SVD took 36911 seconds and DAPL took only 60 seconds, showing that DAPL is several orders of magnitude faster at evaluation time.

## Conclusion

We have described a new method, DAPL, on autoencoder neural network to impute missing data in large genomic datasets. Our results showed that with the modified partial loss it is possible to achieve significant improvement over standard denoising autoencoder. We have simulated multiple missing mechanisms including both missing-at-random cases across a range of missing percentages, and missing-not-at-random cases based on expression level and GC-content. DAPL achieves similar imputation accuracies to the best performing competitor algorithm SVD. We also showed that imputation accuracy is not necessarily compromised by reducing model complexity. This is advantageous for working with large dataset such as in pan-cancer analysis (Cline, et al., 2013; Liu, et al., 2018).

DAPL has the benefit of reducing computational cost especially at evaluation time compared to traditional methods such as SVD and KNN. This is because an autoencoder model can be pre-trained and applied directly to any given new sample to impute missing values, while SVD and KNN methods are not model-based, and hence need to compute all values each time a new sample is given. DAPL provides a strong alternative to traditional methods for estimating missing values in large datasets.

